# Prevalence of loss-of-function alleles does not correlate with lifetime fecundity and other life-history traits in metazoans

**DOI:** 10.1101/224436

**Authors:** Aleksandra V. Bezmenova, Georgii A. Bazykin, Alexey S. Kondrashov

## Abstract

Natural selection is possible only because all species produce more offspring than what is needed to maintain the population. Still, the lifetime number of offspring varies widely across species. One can expect natural selection to be stronger in high-fecundity species. We analyzed the prevalence of loss-of-function alleles in 32 metazoan species and have found that, in contrast to this expectation, the strength of negative selection does not correlate with lifetime fecundity, as well as with other life-history traits. Perhaps, higher random mortality in high-fecundity species negates the effect of increased opportunity for selection.

## Introduction

In the long run, the size of every population that does not go extinct remains approximately constant. Thus, in the course of many generations the geometric mean number of daughters of a female surviving to reproduce must always be one. However, lifetime fecundity of all species is way above this minimum. “There is no exception to the rule that every organic being naturally increases at so high a rate, that if not destroyed, the earth would soon be covered by the progeny of a single pair” (Darwin 1859). In some species, such as elephants and bears, the maximal lifetime fecundity is only ~10, while many others can produce millions of offspring. Of course, to ensure a constant long-term population size, pre-reproductive mortality in a species must be inversely proportional to its average lifetime fecundity.

Production of excessive offspring is a *sine qua non* of natural selection. Indeed, in a species where the maximal lifetime number of daughters is one, any selection would lead to extinction, which could be avoided only if every female produces exactly one daughter. Quantitatively, selection always induces some positive genetic load 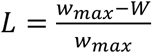, where *w_max_* is the maximal possible fitness and *W* is the mean population fitness (Crow 1970), and, in order for the population size to remain stable, the expected lifetime number of successful daughters of females with the highest fitness must be 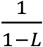. The actual maximal number of daughters of a female must be even larger, due to their pre-reproductive mortality and to random variation of reproduction success among females with the same genotype.

Thus, there is less limitation on the strength of selection in high-fecundity species, which can sacrifice a larger proportion of offspring without going extinct, compared to low-fecundity ones. Therefore, one may expect selection to be stronger in the former. However, this is not necessarily the case, because in high-fecundity species parental investment in an offspring is necessarily low, so that their random mortality is likely to be higher. Thus, while in low-fecundity species mortality of offspring may be mostly due to imperfection of their genotypes and thus lead to selection, in high-fecundity species the bulk of mortality may be irrelevant to selection.

Clearly, the relationship between the maximal fecundity, as well as other life-history traits, of a species and the strength of selection in it needs to be established empirically. With this goal in mind, we compared strengths of negative selection against loss-of-function (LoF) alleles of orthologous genes in 32 metazoan species.

## Methods

We used a large set of transcriptomes published by (Romiguier *et al.* 2014), which consists of sequences of 374 individuals from 76 metazoan species covering 6 phyla (Cnidaria, Annelida, Mollusca, Arthropoda, Echinodermata and Chordata). This dataset also contains information about a number of life-history traits (LHT), such as adult size, body mass, longevity, fecundity, and propagule size. We also collected information about genome sizes of these species, if available (Gregory 2017).

Raw reads were downloaded from the SRA database; SRA accession numbers are listed in Supplemental Table S1, number of sequenced individuals for each species is provided in Supplemental Table S2. They were further trimmed of low-quality positions and sequencing adapters with Trimmomatic software (Bolger *et al.* 2014). Individuals that failed to pass quality control by fastQC (Andrews 2010) after trimming were excluded from further analysis. Reads from all individuals that belong to a species were pooled together and *de novo* assembled into contigs using Trinity (Grabherr *et al.* 2011). Trinity may produce several isoforms of a gene. To exclude minor isoforms, reads were mapped to the assemblies using Bowtie2 (Langmead and Salzberg 2012) and FPKM values were calculated using RSEM program (Li and Dewey 2011). For each gene, we chose an isoform with the highest FPKM value. Open reading frames (ORFs) were predicted using Transdecoder program (with minimum protein length set to 100 amino acids). If more than one ORF were predicted in a contig, the longest ORF was used.

We focused on those genes, hereinafter referred to as core gene, that are present in the list of essential genes of metazoans (Parra *et al.* 2007). A subset of core genes, further referred to as hard-core genes, was obtained by excluding those genes that harbor homozygous LoF mutations. Information about assemblies (number of contigs, N50, mean coverage, alignment rate) and annotated coding sequences (numbers of predicted ORFs and of core genes) are presented in Supplemental Table S2. Species with reads alignment rate less than 70%, number of predicted ORFs less than 5000, or the number of predicted core genes less than 100 were discarded.

For each individual separately, reads were mapped to the reference assembly of the species using Bowtie2. Individuals with mean coverage less than 10x, reads alignment rate less than 80%, number of (at least partially) covered ORFs less than 5000 or the number of (at least partially) covered core genes less than 100 were discarded. SNPs and small indels were called using Samtools mpileup (Li *et al.* 2009) and annotated with Annovar (Wang *et al.* 2010). Only positions with depth more than 5x and mapping and variant calling quality over 20 were considered valid. Stopgain and stoploss substitutions and frameshift indels were assumed to be loss-of-function variants (LoFs).

For each individual, we calculated the proportion of alleles carrying LoF variants among all predicted ORFs and among core genes. Not all coding sequences annotated in reference transcriptomes were sequenced in each individual. The proportion of LoF alleles in an individual was calculated as 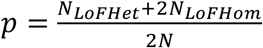, were *N* is the number of ORFs fully sequenced for the individual; *N_LoFHet_* is the number of alleles carrying a heterozygous LoF variant and *N_LoFHom_* is the number of alleles carrying a homozygous LoF variant. This proportion was calculated for all predicted ORFs and for the subsets of core and hard-core genes.

After applying the filters described above, and excluding 8 species of arthropods such as ants, bees and termites whose males are haploid, we ended up with the data set consisting of 32 species, represented by between 1 and 9 individuals, with median 2. LHTs of these species as well as genome sizes and synonymous nucleotide diversities (*π_s_*) obtained from (Romiguier *et al.* 2014) are shown in Supplemental Table S3.

## Results

We recorded the numbers of LoF alleles in genotypes of between 1 and 9 individuals from 32 metazoan species. The proportions of alleles that carry LoF variants among all predicted genes, core genes, and hard-core genes are shown in Supplemental Table S4 (for each individual) and Supplemental Table S5 (mean values for each species). These proportions vary from 0.85% to 5.48% for all genes, from 0% to 5.38% for core genes, and from 0% to 2.12% for hard-core genes, with the means being 2.48%, 1.26%, and 0.30%, respectively. The mean proportion of alleles that carry nonsense substitutions among all genes in all species was 0.47%, while the mean proportion of alleles that carry frameshift indels was 1.78% (Table 1).

**Table 1.** Mean species proportions of LoF alleles among all genes.

|  | Min | Max | Mean |
| --- | --- | --- | --- |
| **All LoF variants** | 0.85% | 5.48% | 2.48% |
| **Nonsense** | 0.071% | 1.03% | 0.47% |
| **Frameshift** | 0.040% | 4.27% | 1.78% |
| **Stoploss** | 0.00% | 0.68% | 0.13% |

We related the mean proportion of LoF alleles in a species to its lifetime fecundity (Figure 1) and other life-history traits, as well as to genome size and *π_S_* (Figure 2). This proportion shows no significant correlation with any of the traits. No correlations were also observed when different types of LoF variants were considered separately (Supplemental Figure S1 and Supplemental Figure S2).

**Figure 1.**
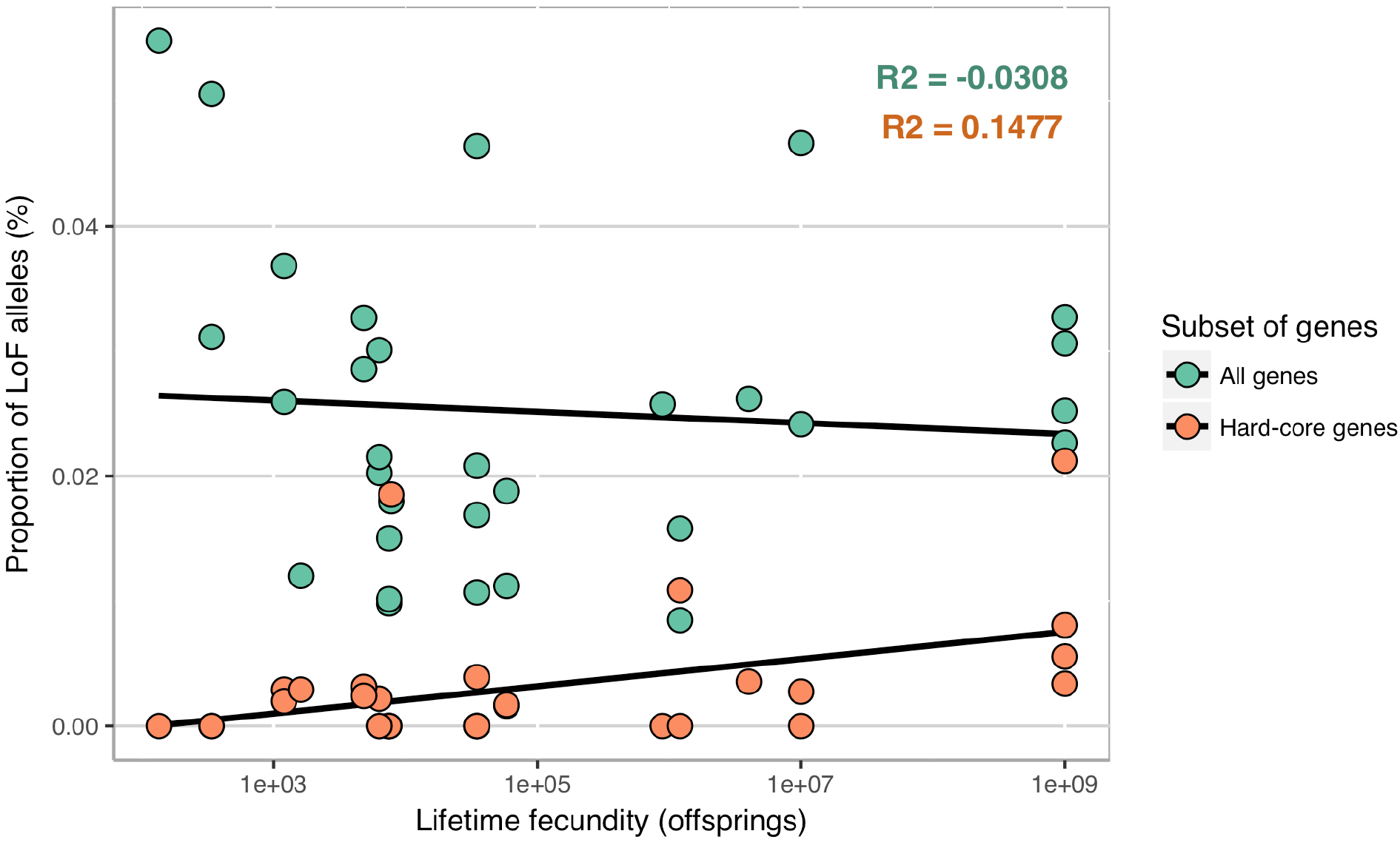
The mean proportions of LoF alleles against lifetime fecundity in all (green) and in hard-core genes (orange) for each species. Spearman’s correlation coefficients are shown.

**Figure 2.**
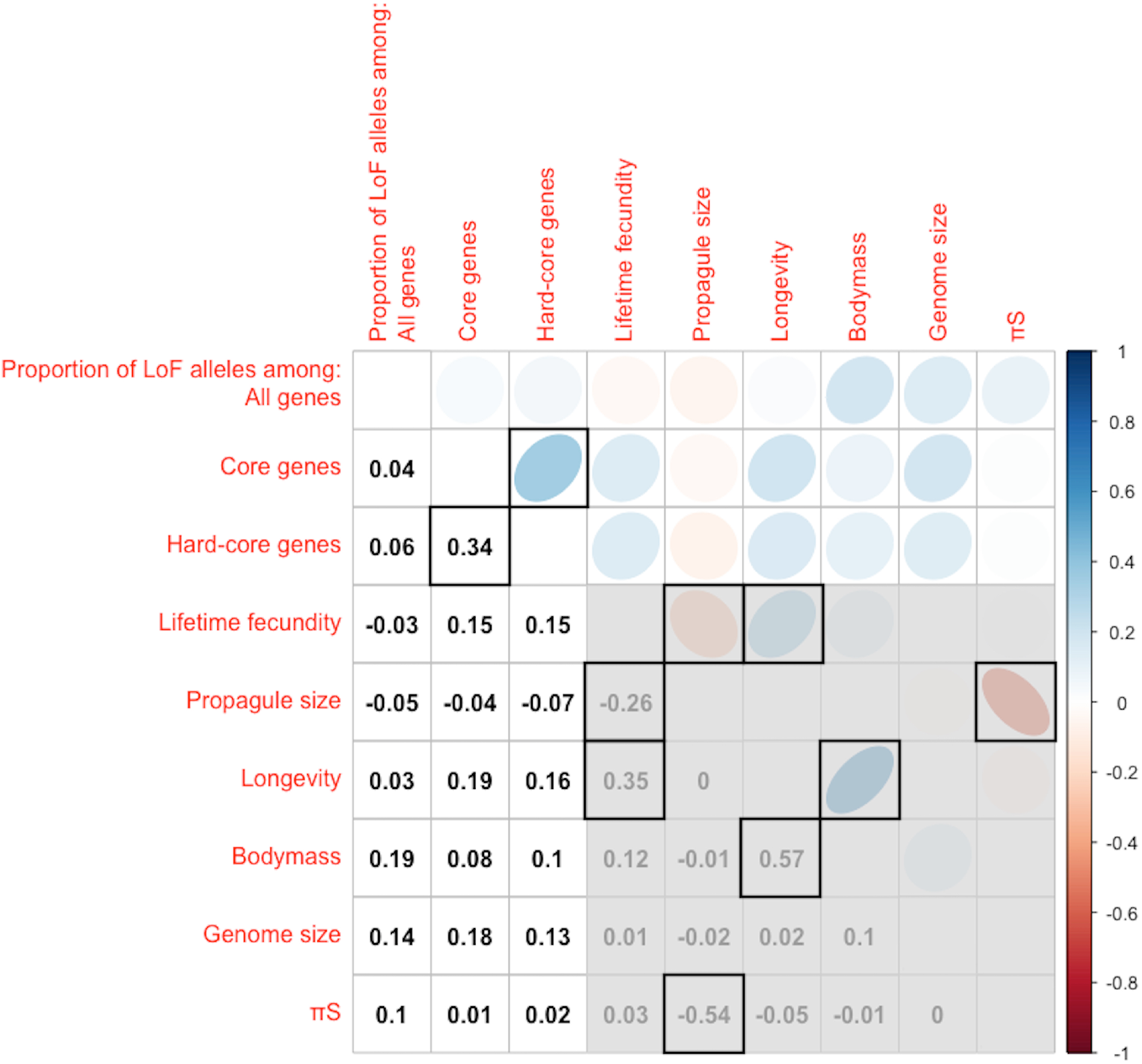
Correlations between mean proportions of LoF alleles among all, core and hard-core genes and life-history traits. Blue indicates a positive relationship, and red, a negative relationship; colour intensity is proportional to Spearman’s correlation coefficients, which are also presented below the diagonal. Correlations that are significant after the BH (Benjamini and Hochberg) correction for multiple testing are framed.

## Discussion

We investigated the strength of negative selection across a wide variety of metazoan species. This strength was assayed through the prevalence of LoF alleles of essential genes in genotypes of individuals. Frequencies of such alleles are generally quite low, and the data on recessive lethals in *Drosophila* populations suggest that coefficients of selection against them, in the heterozygous state, are ~1% (Simmons and Crow 1977). Thus, the frequencies of such alleles are likely to be close to the deterministic mutation-selection equilibrium even in small natural populations (Cassa *et al.* 2017), which almost always have *N_e_* >= 10^4^. In other words, the prevalence of such LoF alleles should be essentially independent of the effective sizes of natural populations. Indeed, in great apes the prevalence of LoF alleles does not depend on the effective population size (de Valles-Ibáñez *et al.* 2016). Our analysis also found no correlation between the proportion of genes carrying LoF mutations and π_S_, an estimator of the effective population size. From this perspective, LoF alleles of important genes are radically different from missense mutations of all protein-coding genes, which are more prevalent in species with low effective population sizes due to inefficient selection against slightly deleterious mutations (Popadin *et al.* 2007).

The mean proportion of all genes carrying LoF variants across all species was 2.5%, comparable to those reported for primates, from ~0.7% in *Homo sapiens* to ~2.2% in *Pongo abelii* (de Valles-Ibáñez *et al.* 2016). The proportion of frameshift indels exceeded the proportion of nonsense substitutions by a factor of ~4.6, which is close to the values obtained in (de Valles-Ibáñez *et al.* 2016), 1.7 – 4.7.

We observed no significant correlations between the prevalence of LoF alleles in all, core, or hard-core genes and lifetime fecundity or any other life-history trait of a species. This suggests that the random mortality in highly prolific species may negate a higher opportunity for natural selection.

Of course, the prevalence of LoF variants must be proportional to the mutation rate. Could this fact mask the positive dependence of the strength of negative selection on the lifetime fecundity? This seems to be unlikely. Indeed, in order to explain our result in this way, one needs to assume that high-fecundity species have higher mutation rates. However, no data support this hypothesis. In fact, there are week correlations of the opposite sign, as high-fecundity species tend to have higher *N_e_* (Figure 2), and species with higher *N_e_* tend to have higher mutation rates (Lynch *et al.* 2016). We also observe no strong correlation between the prevalence of LoFs and π_S_, which should also depend linearly on the mutation rate.

Our analysis should not be confounded with the studies of the impact of random drift on the action of weak selection with |s| ~ 1/*N_e_* (Kimura 1983). The efficiency of weak negative selection has been shown to decline in small populations, where more polymorphisms become effectively neutral (Popadin *et al.* 2007; Nikolaev *et al.* 2007). In contrast, negative selection against the majority of even heterozygous LoF variants is sufficiently strong (Simmons and Crow 1977; Cassa *et al.* 2017) to make their dynamics essentially independent of the random drift even in the smallest natural populations ((de Valles-Ibáñez *et al.* 2016) and Figure 2).

Thus, it appears that a heterozygous LoF variant of a particular gene causes the same reduction of fitness in species with drastically different lifetime fecundities and opportunities for selection. This invariance is puzzling. Could it be a consequence of the evolutionary optimization of fecundity and other LHTs? If all species possess the values of LHTs that lead to the highest fitness, given their particular constraints, this may lead to the strongest possible negative selection. Still, it is not clear why the strongest possible selection turns out to be equally strong in a cod and elephants.

## Availability of data and materials

Raw data is available in the SRA database, accession numbers are listed in Supplementary file 3, Table S1. Assembled transcriptomes and other data are available upon request.

## Acknowledgment

This work was supported by the Russian Science Foundation, grant № 1614-10173.

## Authors’ contributions

AB carried out the data analyses, participated in the design of the study and drafted the manuscript; GB participated in the design of the study and helped draft the manuscript; AK conceived of the study and helped draft the manuscript. All authors gave final approval for publication.

